# Effects of Human Sex Chromosome Dosage on Spatial Chromosome Organization

**DOI:** 10.1101/346742

**Authors:** Ziad Jowhar, Sigal Shachar, Prabhakar R. Gudla, Darawalee Wangsa, Erin Torres, Jill L. Russ, Gianluca Pegoraro, Thomas Ried, Armin Raznahan, Tom Misteli

## Abstract

Sex chromosome aneuploidies (SCAs) are common genetic syndromes characterized by the presence of an aberrant number of X and Y chromosomes due to meiotic defects. These conditions impact structure and function of diverse tissues, but the proximal effects of SCA on genome organization are unknown. Here, to determine the consequences of SCAs on global genome organization, we have analyzed multiple architectural features of chromosome organization in a comprehensive set of primary cells from SCA patients with various ratios of X and Y chromosomes by use of imaging-based high-throughput Chromosome Territory Mapping (HiCTMap). We find that X chromosome supernumeracy does not affect the size, volume or nuclear position of the Y chromosome or an autosomal chromosome. In contrast, the active X chromosome undergoes architectural changes as a function of increasing X copy number, as measured by a decrease in size and an increase in circularity, which is indicative of chromatin compaction. With Y chromosome supernumeracy, Y chromosome size is reduced suggesting higher chromatin condensation. The radial positioning of chromosomes is unaffected in SCA karyotypes. Taken together, these observations document changes in genome architecture in response to alterations in sex chromosome numbers and point to trans-effects of dosage compensation on chromosome organization.

## Introduction

Sex chromosome aneuploidies (SCAs) are a class of genetic human syndromes that arise by to the presence of an aberrant number of X and/or Y chromosomes beyond the typical XX for females and XY for males (Disteche, 2012). Commonly studied SCAs include Turner syndrome (X-monosomy in females (XO), and Klinefelter syndrome, which arises due to presence of a supernumerary X-chromosome in males (XXY karyotype), although other variations include XYY, XXX, and XXYY syndrome. These conditions are collectively common (prevalence 2.5/1000), and associated with neurological, endocrine, immune and metabolic phenotypes (Nielsen and Wohlert, 1991; Linden *et al.*, 1995; Bojesen *et al.*, 2003; Goswami *et al.*, 2003; Simpson *et al.*, 2003; Stochholm *et al.*, 2006; Chen *et al.*, 2012; Link *et al.*, 2013; Seminog *et al.*, 2015; Mankiw *et al.*, 2017). The downstream phenotypic consequences of SCAs presumably arise from the impact of atypical sex chromosome counts on genome function, but genomic features of SCA remain poorly understood in humans (Bermejo-Alvarez *et al.*, 2010; Arnold, 2012; Hughes and Rozen, 2012; Belling *et al.*, 2017).

In order to maintain cellular function, dosage compensation between males and females in mammals is achieved by silencing most of the genes in one of the two X chromosomes through a mechanism known as X chromosome inactivation (XCI), thus generating two functionally different X chromosomes, the inactive X (Xi) and active X (Xa) (Clemson *et al.*, 1996; Avner and Heard, 2001). The initiation of XCI requires the non-coding *Xist* (X-inactive-specific transcript) RNA, a 17 kb-long untranslated transcript, which decorates Xi territories in cis (Brown *et al.*, 1991; Clemson *et al.*, 1996). *Xist* recruits various epigenetic factors that induce heterochromatinization of the X chromosome, including histone methylation (i.e., H3K27me3) and deacetylation (H4 hypoacetylation) via PRC1/2 and histone deacetylases (HDACs), respectively, repressive histones (macroH2A), and incorporation of DNA methylation (CpG islands) by various DNA methyltransferases (DNMTs) (Plath *et al.*, 2002; de Napoles *et al.*, 2004; Chaumeil *et al.*, 2006; Engreitz *et al.*, 2013).

Given the presence of extra copies of sex chromosomes in SCAs, an unresolved question is whether the abnormal number of sex chromosomes leads to changes in the organization or spatial location of chromosomes within the nucleus. Chromosomes are spatially confined to discreet nuclear volumes, each referred to as chromosome territory (CT) (Schardin *et al.*, 1985; Cremer *et al.*, 1993; Kurz *et al.*, 1996; Meaburn and Misteli, 2007) and they have characteristic, heterogeneous, but non-random distributions within the 3D space of the nucleus (Parada *et al.*, 2004b; Misteli, 2007) (Borden and Manuelidis, 1988; Croft *et al.*, 1999; Bartova *et al.*, 2002; Parada *et al.*, 2004a). Interestingly, the Xi forms a distinct, condensed and spherical structure (Barr body) and has a more peripheral position within the nucleus compared to the Xa chromosome (Chadwick and Willard, 2003; Bolzer *et al.*, 2005; Teller *et al.*, 2011).

In this study, we take advantage of a unique, comprehensive set of cells from individuals with various SCAs including XO, XXX, XXXX, XXY, XYY, XXYY and XXXXY (Table 1) to probe the effect of sex chromosome supernumeracy on chromosome organization. Our studies are enabled by the recent development of HiCTMap, a 384-well plate based chromosome paint FISH imaging method (Jowhar *et al.*, 2018b) to characterize the size, shape and positioning of chromosomes in very large numbers of cells at high-throughput. We find that in SCA the X chromosome undergoes architectural changes as a function of X copy number indicated by a decrease in size and increase in circularity, suggesting chromatin compaction, while the Y chromosome decreases in size as a function of Y copy number. All chromosomes studied maintain canonical spatial positioning regardless of sex chromosome numbers. Our results provide insight into the structural consequences of changes in genome organization induced by sex chromosome supernumeracy.

**Table 1.** Primary Skin Fibroblast Cell Line Description and Origin.

| **Karyotype** | **Patient Number** | **Sex** | **Age** | **Tissue Origin** |
| --- | --- | --- | --- | --- |
| XO | FA | F | 21 | Skin punch biopsy |
| XX | FA | F | 22 | Skin punch biopsy |
| XXX | FA | F | 14 | Skin punch biopsy |
| XXXX | FA | F | 24 | Skin punch biopsy |
| XXXXY | FA | M | 16 | Skin punch biopsy |
| XXY | FA | M | 16 | Skin punch biopsy |
| XXYY | FA | M | 21 | Skin punch biopsy |
| XY | FA | M | 17 | Skin punch biopsy |
| XYY | FA | M | 24 | Skin punch biopsy |
| XO | FB | F | 13 | Skin punch biopsy |
| XX | FB | F | 18 | Skin punch biopsy |
| XXX | FB | F | 14 | Skin punch biopsy |
| XXXX | FB | F | 11 | Skin punch biopsy |
| XXXXY | FB | M | 13 | Skin punch biopsy |
| XXY | FB | M | 19 | Skin punch biopsy |
| XXYY | FB | M | 12 | Skin punch biopsy |
| XY | FB | M | 8 | Skin punch biopsy |
| XYY | FB | M | 21 | Skin punch biopsy |
| XO | FC | F | 14 | Skin punch biopsy |
| XX | FC | F | 17 | Skin punch biopsy |
| XXX | FC | F | 16 | Skin punch biopsy |
| XXXX | FC | F | 12 | Skin punch biopsy |
| XXXXY | FC | M | 16 | Skin punch biopsy |
| XXY | FC | M | 16 | Skin punch biopsy |
| XXYY | FC | M | 13 | Skin punch biopsy |
| XY | FC | M | 25 | Skin punch biopsy |
| XYY | FC | M | 19 | Skin punch biopsy |

Description of patient derived skin fibroblast cell lines used for experiments. Karyotypes, cell line name, ages, and tissue origins are indicated.

## Results

### Detection and analysis of chromosome territories using HiCTMap in nuclei from SCA patients

In order to identify the effects of extra copies of either X and/or Y chromosomes on spatial genome organization, we applied the previously described high-throughput chromosome territory mapping (HiCTMap) method (Jowhar et al 2018). HiCTMap is a 384-well plate based imaging method for high-throughput, quantitative structural analysis and mapping of the spatial location of chromosome territories (CTs) in the mammalian cell nucleus (Jowhar et al 2018). HiCTMap allows for systematic, high-resolution imaging of thousands of nuclei in hundreds of samples and it can accurately measure structural features of chromosomes such as size, shape and spatial position at the single chromosome level (Jowhar et al. 2018). We applied this pipeline to a comprehensive set of patient-derived primary cell lines which were acquired via skin biopsy from seven SCA patients (XO, XXX, XXXX, XXXXY, XXY, XXYY, XYY) and normal male (XY) and female (XX) controls (Figure 1A, Table S1). Skin fibroblast cell lines from 3 patients per karyotype that ranged from ages 8-25 were analyzed (Table S1). Cells were routinely cultured for no more than 3-7 passages prior to analysis to minimize the possibility of genomic changes and their karyotype was confirmed by spectral karyotyping (SKY) (data not shown). Normal cell growth was observed as assessed the proliferation marker Ki-67 (Supplementary Figure S1) and normal cell cycle profiles were observed at various cell densities (Supplementary Figure S2). Chromosomes 18, X, and Y were routinely detected by HiCTMap in all cell lines (Figure 1, A and B). For image analysis, we used a modified KNIME based image analysis workflows (Ronneberger *et al.*, 2015; Gudla *et al.*, 2017; Jowhar *et al.*, 2018a) (see Materials and Methods). These workflows extract various geometric and intensity-related features of the segmented CTs, including chromosome area and circularity, as well as their spatial position in the nucleus (Figure 1A). We typically analyzed a minimum of 1,000 cells in at least three experimental replicates per karyotype per experiment.

**Figure 1.**
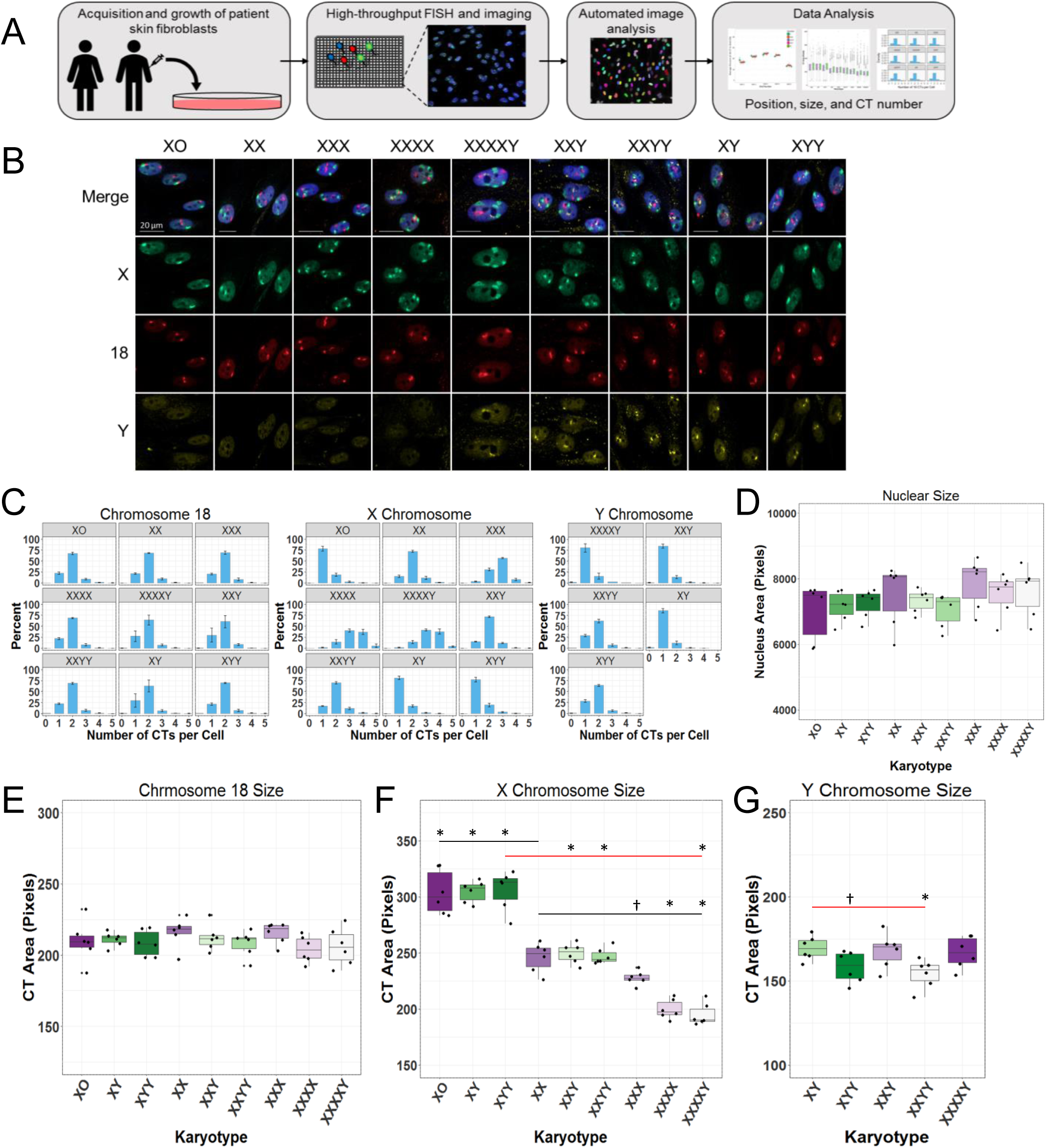
HiCTMap Detection in SCA Nuclei. A) HiCTMap outline. Cells are acquired from patients via skin biopsy and expanded in cell culture. Cells are plated in 384-well imaging plates and DNA FISH is carried out using chromosome paints, followed by automated image acquisition using high-throughput microscopy. Image analysis by KNIME segments the nuclei, detects CTs in three channels, and measures Chromosome position, size, and CT number. These features are then plotted using the R software. B) Representative maximal projections of CT in a set of SCA skin fibroblasts. Cells are stained with DAPI (Blue-408) and chromosome paint probes X-Green (Alexa488), 18-Red (Dy505), Y-Red (Dy505), and Y-FarRed (Dy651). Scale bar: 20 μm. C) Histograms showing the number of CTs detected per nucleus in SCA and normal male and female cells. Bars represent mean of 3 biological replicates, each replicate contains ~1500 cells analyzed per karyotype. D) Nuclear area of all karyotypes. E-G) CT area of chromosomes 18 (E), X (F) and Y (G). Boxes represent the 25th, 50th (median) and 75th percentile of the distributions and whiskers extend to 1.5X the inter-quantile range (IQR). Each dot represents a replicate which contained ~1,000 nuclei per karyotype. The red bar compares normal males to male SCA karyotypes. The black bar compares normal female to all karyotypes. F) *: p-value = 0.0001, †: p-value = 0.07. G) *: p-value < 0.05, †: p-value = 0.12. Statistics calculated using one-way ANOVA, and only significant differences are denoted.

We first sought to verify the aneuploidy status of the multiple karyotypes using HiCTMap. The number of X, Y, and as a control, chromosome 18 CTs was determined for each karyotype. The control chromosome 18 was detected in two copies in 69% of cells when averaged across all karyotypes (Figure 1C). The expected number of X and Y chromosomes was detected in 40 to 80% of cells depending on karyotype (Figure 1C). As expected and previously described (Jowhar *et al.*, 2018b), the number of individually distinguishable X chromosomes is lower in multi-X karyotypes (XXX, XXXX, XXXXY) than the actual number due to clustering of multiple homologue chromosomes which consequently appear as a single, enlarged CT (Jowhar *et al.*, 2018b). Such clustering was observed in 20% to 60% of cells depending on karyotype with the extent of clustering correlating with the number of X chromosomes (Figure 1C). In these cases, the reduced CT number was not a limitation of the image analysis segmentation algorithm since visual inspection was equally unable to resolve the clustered chromosomes. For all further analyses, only nuclei that contained the expected number of chromosomes per karyotype were used.

### Chromosome size upon increased X or Y copy number

Addition of extra copies of sex chromosomes could potentially lead to an increase in nuclear volume. To test whether nuclear size changes with increasing amount of DNA in SCA karyotypes, we measured the mean nuclear area of all SCA karyotypes (see Materials and Methods). We find that the nuclear area of all SCA karyotypes does not show a statistically significant difference, irrespective of the number of X or Y chromosomes when compared to the corresponding normal XX or XY controls (One-way ANOVA, p-value > 0.05 for all comparisons, Figure 1D). Since nuclear size does not significantly change, we did not normalize CT size to nuclear area in further analyses.

We next measured the CT size of control autosome 18 and sex chromosomes X and Y in all karyotypes (Figure 1, E-G). We find that the median area of autosome 18 is similar between different karyotypes with no statistically significant difference between karyotypes when compared to their corresponding normal control karyotypes XX or XY (p-value > 0.05, Figure 1E) (mean +/- SD, range of XXXX=204 +/- 9 to XX=217 +/- 10, p- value > 0.05). In contrast, the range of the median X chromosome area decreased with increasing X chromosome copy number (Figure 1F) from 313 +/- 18 pixels in XYY to 190 +/- 10 in XXXXY; p-value < 0.05), as would be expected from compaction of inactivated X-chromosomes. The medianCT area was increased by ^~^25% between monosomy X karyotypes (XO, XY, XYY) and normal XX females (p-value = 0.0001). In tetrasomy X karyotypes (XXXX, XXXXY), the median CT area decreases by 20% and 35% compared to XX or XY, respectively (p-value = 0.0001; Figure 1F). These changes in mean X chromosome area with increasing X copy numbers are in line with the expectation of X inactivation of any supernumerary X chromosomes.

The median area of the Y chromosome did not differ between single Y karyotypes compared to normal XY (p-value > 0.9). However, unexpectedly, we see a decrease in median Y CT area in XXYY and in XYY cells compared to XY (XY vs XXYY: p-value = 0.02, XY vs XYY: p-value = 0.12) (Figure 1G). This decrease may be suggestive of a higher-order regulatory mechanism to compensate for the increase Y.

### X chromosome inactivation status in nuclei from SCA patients

It is well established that in cells from normal XX females, one X chromosome undergoes inactivation, and it is expected that with increasing X chromosome copy numbers the number of Xi chromosomes should increase so that only one Xa is maintained in non-cancerous polysomic X karyotypes (Lyon, 1961; Therman *et al.*, 1980; Sarto *et al.*, 1987). We expanded HiCTMap to combine DNA and RNA FISH to detect both X CT using chromosome paints and *Xist* RNA using RNA FISH probes (see Methods & Material). Xa- and Xi-territories were clearly distinguishable based on co-localization of *XIST* transcripts on the Xi CT (Figure 2). We find that in nuclei from all karyotypes only one X CT remains active, whereas all other X CTs undergo inactivation as determined by *Xist* staining (Figure 2A). Generally, the number of Xist foci correlated better with the expected number of X chromosomes than the X CT itself, since Xist covers a small, compact area that is more easily resolved both visually and with computational methods. When comparing three patient cell lines with multi X karyotypes, we detected the correct number of *Xist* stained X chromosomes in similar number of cells per karyotype, which is indicative of little inter-patient variability (Figure 2B). Furthermore, the low variance in XIST focus count within each karyotype group indicated low cellular heterogeneity in X-inactivation status. No visible Xist foci were detected in monosomy X karyotypes (Supplementary Figure S3, A and B). For further analyses, only nuclei containing the expected number of *Xist* foci were included.

**Figure 2.**
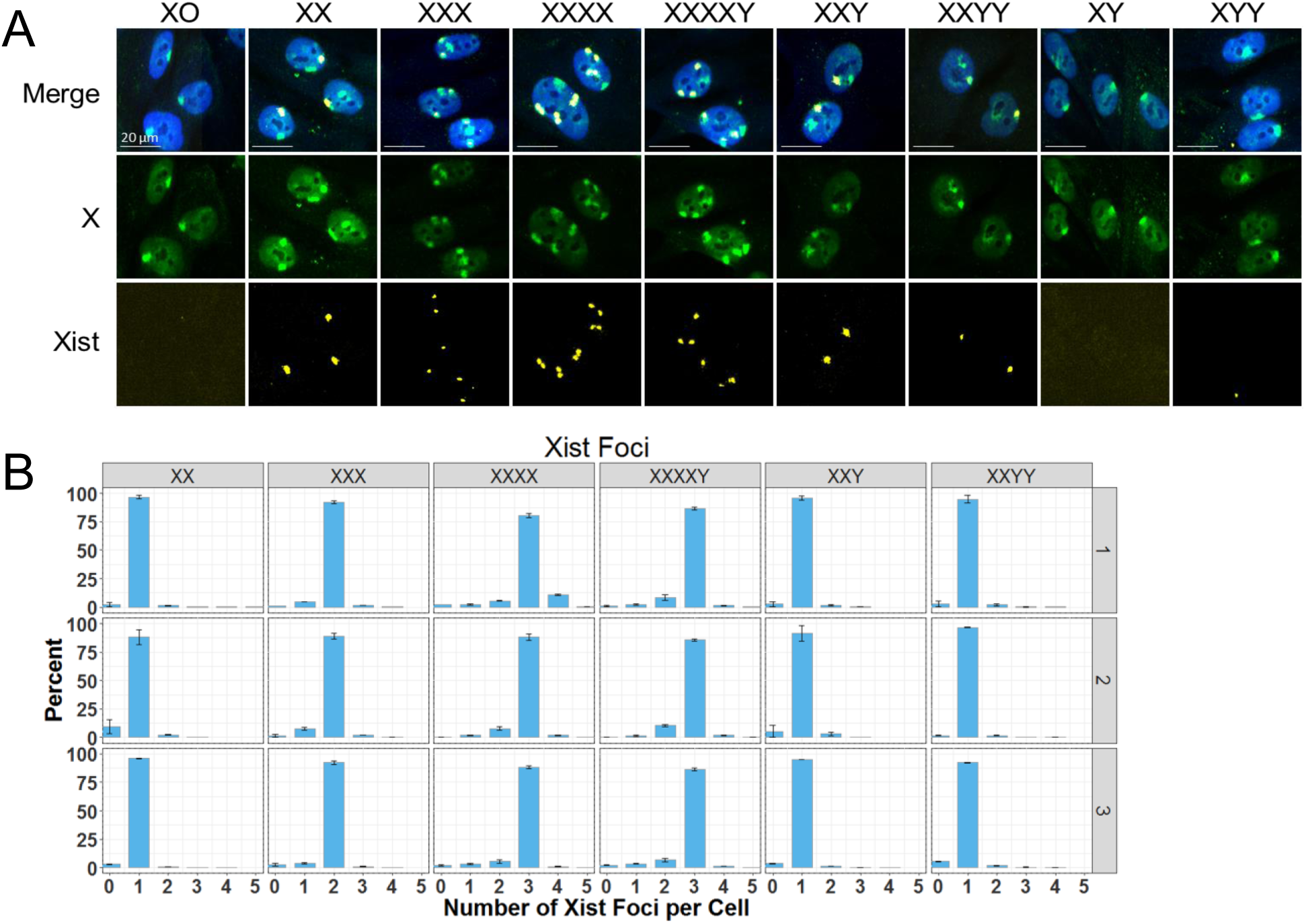
Xist RNA in SCA Nuclei. A) Representative maximal projection images of combined X chromosome DNA FISH and *Xist* RNA FISH. Cells are stained with DAPI (Blue-408) and X chromosome paint probes and Stellaris Xist Design Ready probe. Scale bar: 20 μm. B) Number of Xist foci detected per nucleus in polyploid X nuclei. Error bars represent ± SD. Bars represent the means between 3 biological replicates, each replicate contains approxiametly1, 000 nuclei analyzed per karyotype.

### Changes in X chromosome size and shape with increasing X dosage

To determine whether the architecture of the X chromosome changes in SCAs, we measured structural features of X CT including size and shape of both Xa and Xi (Figure 3). As expected, we observe a statistically significant difference between the mean Xa and Xi area in all SCA karyotypes (Xa>Xi, Two-way ANOVA, p < 0.05 for all comparisons). However, the difference in size between Xa and Xi decreased with increased X dosage (Figure 3A). This was due to a marked reduction of the median size of the Xa CT with increasing X dosage (mean +/- SD, XX = 282 +/- 20, XXX = 259 +/- 8, XXXX = 229 +/- 12, XXXXY = 222 +/- 14, XXY = 277 +/- 4, & XXYY = 276 +/- 9 pixels) (Fig 3A, red dots). In tetrasomy X karyotypes, the mean Xa CT size decreases by as much as 20% to 30% compared to XX or XY, respectively (p-values = 0.0001). Similarly, the Xa of monosomy X karyotypes is larger than the Xa in XX cells by 12% (p-value < 0.0022). The Xi chromosome also decreased in size with increasing X copy number, although to a smaller extent (XX = 214 +/- 13, XXX = 211 +/- 6, XXXX = 189 +/- 10, XXXXY = 184 +/- 11, in XXY = 225 +/- 17, & XXYY = 224 +/- 13 pixels) (Figure 3A, blue dots). The size of Xi in XXXX and XXXXY was reduced by 12% and 15%, respectively, when compared to XX, representing the normal X karyotype and containing a single Xi (p-value < 0.01). The size of Xi in disomy X karyotypes did not change significantly (p-value > 0.05). Additionally, trisomy X exhibits a 3% decrease in mean Xi CT area compared to XX, but this difference was not statistically significant (p-value > 0.05).

**Figure 3.**
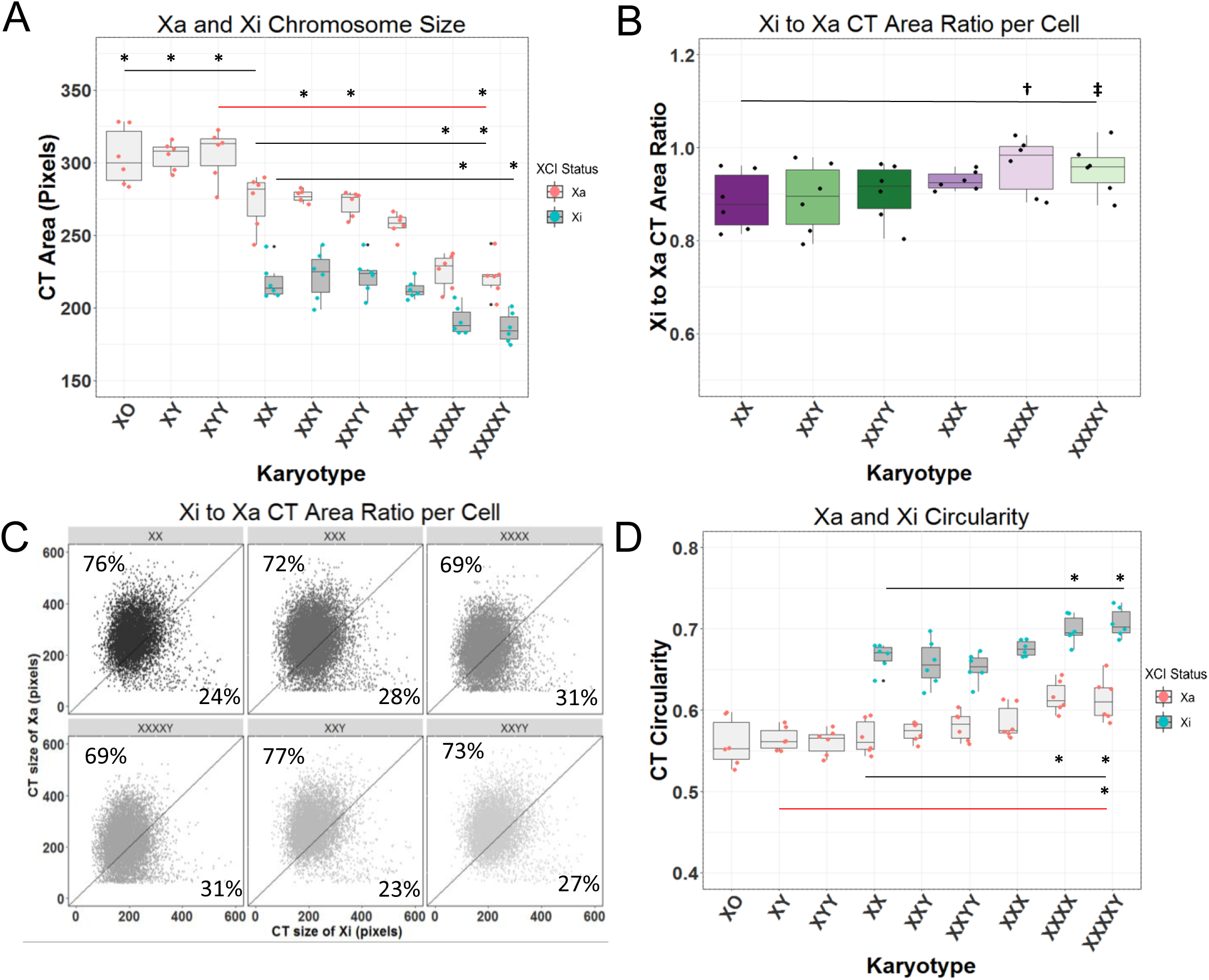
Chromosome Xa and Xi Structural Characterization in SCA Nuclei. A) CT area of Xa and Xi in SCAs and normal male and female nuclei. Red dots represent Xa CTs and blue dots represent Xi CTs. *: p-value < 0.003. B) Xi to Xa CT area ratio on a per cell basis of polyploid X karyotypes. Each spot represents the mean of XXX replicates †: p-value = 0. 12, ‡: p-value = 0.19. C) Xa CT area vs Xi CT area on a per Xa basis of polyploid-X karyotypes. Each dot represents a single nucleus. Percentages represent the portion of ratios in which Xa is larger than Xi D) CT circularity of Xa and Xi in SCAs and normal male and female nuclei. Circularity is measured as perimeter^2^/area *: p-value < 0.05. A, B, D) All box plots show the 25th, 50th (median) and 75th percentile of the distributions and whiskers extend up to 1.5X of the inter-quantile range. Each dot represents a replicate each containing ~1,000 nuclei per karyotype. *: p-value < 0.05. The red bar compares normal males to male SCA karyotypes. The black bar compares normal female to all karyotypes. Statistics calculated using one-way ANOVA, and only significant differences are denoted.

The observation of a marked change in size of X CT was based on measurement of the mean X CT size in a population of cells. To determine whether the extent of X CT size reduction per cell is different between different karyotypes, we measured the ratio between Xi and Xa size on a per-cell basis to determine whether in each individual cell, all X CTs are reduced to the same extent (Figure 3B). The median Xi/ Xa ratio in normal XX cells ranged from 0.88 +/- 0.06 in XX cells to XXXX = 0.98 +/- 0.06 in XXXX cells, suggesting that the Xa is decreased more drastically with increasing X copy number than the Xi (Figure 3B). To verify, we plotted the size of Xa CT vs the size of Xi CT on a per-cell basis (Figure 3C) and find that in 69% − 77% of cells, Xa is larger than Xi (Figure 3C). These observations demonstrate considerable variability in Xi and Xa size in individual cells and a size effect on both X chromosomes with increased X copy number.

Since we detected significant size changes in the X chromosome with increasing X copy number, we wanted to further characterize the structure of the Xa and Xi CTs in SCA karyotypes. We specifically measured the circularity of the chromosomes (defined as area/perimeter^2^), as previous reports demonstrated that increased roundness or circularity of chromosomes is associated to compaction and heterochromatinization of the inactivated X chromosome (Bischoff *et al.*, 1993; Eils *et al.*, 1996; Clemson *et al.*, 2006; Teller *et al.*, 2011). As with size, the circularity of Xa chromosomes is X copy number dependent with an increase in mean Xa circularity from 0.57 +/- 0.02 in XX to 0.61 +/- 0.02 in XXXXY, and demonstrate a stiditically significant difference when compared to XX or XY (p-value < 0.005; Fig 3D, red dots). The mean Xi CT circularity, like size, increased from 0.67 +/- 0.02 in XX to 0.7 +/- 0.02 in XXXXY (p-value < 0.02; Figure 3D, blue dots). This change in circularity is unique to the X chromosome as we do not see a change in the mean circularity of chromosomes 18 or Y (mean +/- SD, 0.66 +/- 0.02) in all SCA karyotypes with varying X copy number on average (Supplementary Figure S4). The circularity measurements of chromosomes 18 and Y, which are both relatively small in size, gene poor, and lowly expressed comparable to the Xi CT, which is transcriptionally inactive and highly compacted. In contrast, circularity of the transcriptionally active X chromosome is lower, suggesting that circularity measurements are a good estimate for the overall chromatin compaction state of chromosomes. Taken together, the increases in circularity observed for both the Xa and Xi with X copy number are suggestive of changes in the chromatin structure via an increase in heterochromatin in the presence of supernumerary X chromosomes.

### Spatial positioning of chromosomes X, Y and 18 in SCA nuclei

Due to the nonrandom organization of the genome and observations that chromosome positioning may affect cellular function, we sought to test whether additional sex chromosomes affect the location of chromosome X, Y and 18 in the nucleus. Using HiCTMap as previously described (Jowhar *et al.*, 2018b) we mapped the radial position relative to the center of the nucleus of CTs 18, X, and Y in all SCA karyotypes. For analysis, nuclei were divided into five equidistant shells, and the percentage of each CT in each shell was calculated (Croft *et al.*, 1999; Tanabe *et al.*, 2002; Bolzer *et al.*, 2005). We find that the radial position of chromosome 18 does not change between all SCA karyotypes with 12-24% of 18 CTs occupying each shell as shown previously in fibroblasts (Bolzer *et al.*, 2005) (Figure 4A). As previously reported for XY, the Y chromosome was preferentially found in the interior of the nucleus with no significant difference between the various karyotypes (Figure 4B). The Y CT occupies shells 3 and 4 with the highest percentage in all male SCA karyotypes (Figure 4B).

**Figure 4.**
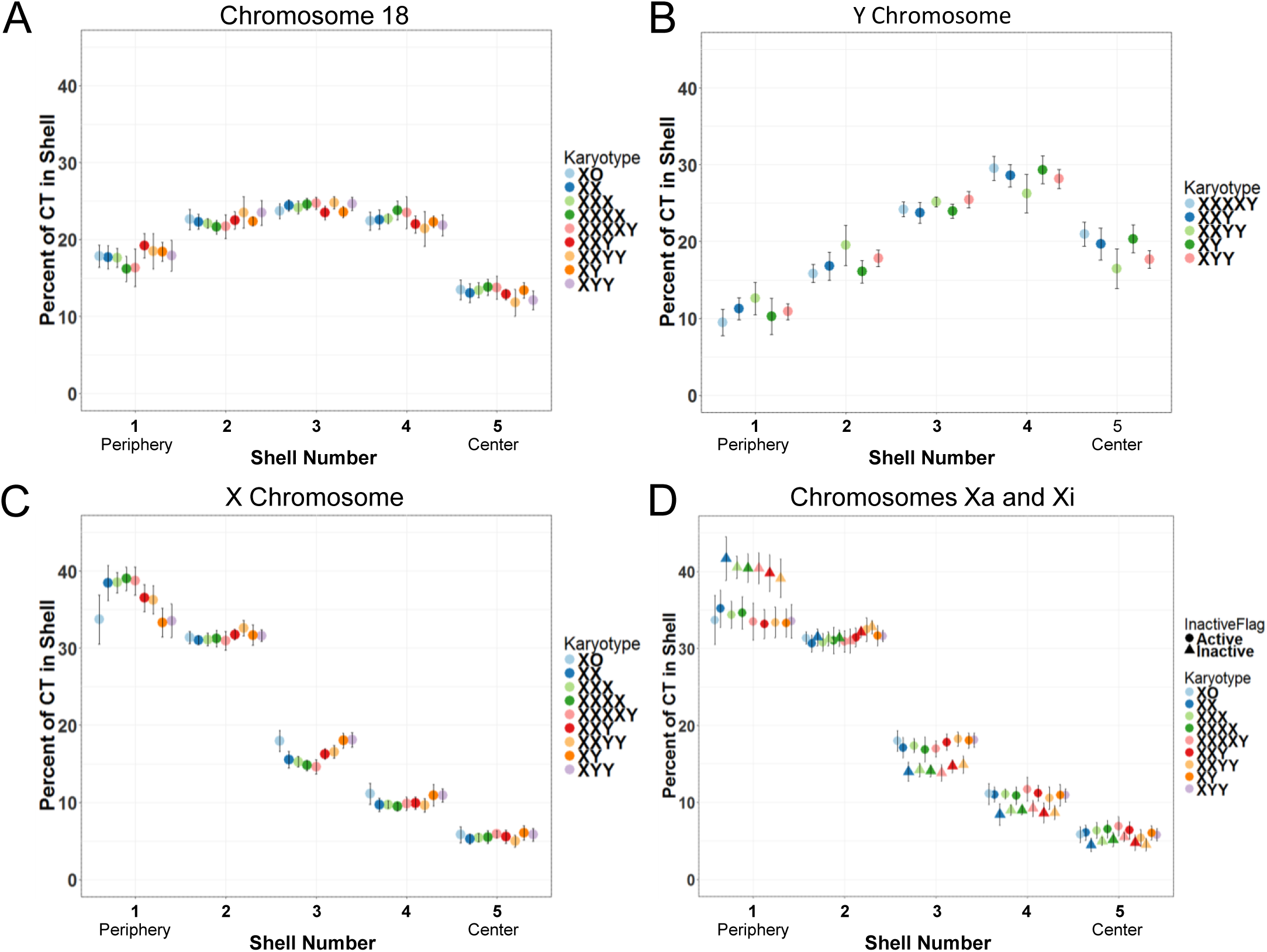
Radial Positioning of Chromosomes 18, X, and Y in SCA Nuclei. A) Distribution of the percentage of chromosome 18 in 5 concentric equidistant nuclear shells. Each spot represents the average of 6 replicates, (3 biological replicates, 2 experimental repeats each), each replicate contained a of ~1,000 nuclei. B) Distribution of the percentage of chromosome Y in 5 concentric equidistant nuclear shells. C) Distribution of the percentage of chromosome X in 5 concentric equidistant nuclear shells. D) Distribution of the percentage of chromosome Xa and Xi in 5 concentric equidistant nuclear shells. Triangles represent Xi CTs and circles represent Xa CTs. Error bars represent ± SD.

The X chromosome shows a strong peripheral positioning, as reported previously (Belmont *et al.*, 1986; Nagele *et al.*, 1999; Bolzer *et al.*, 2005; Petrova *et al.*, 2007) (Figure 4C). With increasing X copy number, the positioning of the entire population of X chromosome becomes slightly more peripheral, due to the presence of the additional Xi chromosomes, which tend to be more peripheral than the Xa (Figure 4C) (Belmont *et al.*, 1986; Chen *et al.*, 2016). When positioning is measured individually for the Xa or Xi, both chromosomes maintain a strong peripheral positioning, with Xi exhibiting a more peripheral location compared to Xa in all multi-X karyotypes with a concurrent loss of Xi in the interior (Figure 4D; p-value < 0.05). We conclude that SCAs does not affect positioning of the Xa or Xi.

## Discussion

We have used HiCTMap, a high-throughput imaging pipeline for the structural characterization and mapping of chromosome positions, to probe the effect of sex chromosome aneuploidy on genome architecture. The ability to analyze a large number of cells by high-throughput imaging enabled us to quantitatively determine the structural features and radial position of chromosomes in a unique set of primary cell lines from 27 SCA patients representing distinct karyotypes.

We find that SCAs do not induce changes in the size of the nucleus or the size of autosomal chromosome 18. However, we find that sex chromosome Y shows a decrease in its chromosome size with an increase in Y copy number, but not with increased X copy number. Similarly, we see changes in sex chromosome X size with the increase in X copy number, but not Y copy number. While the latter observation is explained by the well-established X inactivation process used to compensate for the increase number of X chromosomes (Lyon, 1961; Boumil and Lee, 2001; Willard, 2014), the mechanism for the reduction of Y size is unknown. Chromosome Y is small, gene poor, and largely heterochromatic and the observed size change may be due to a mechanism by which cells attempt to compensate for the increase in gene dosage via changes in chromatin structure much like XCI, but to a lower extent. The effect on the Y chromosome is likely mediated by epigenetic mechanisms and may be necessary for maintaining cellular viability given the known capacity of some Y-linked genes, like SRY, to regulate the expression of autosomal and X-linked genes whose increased expression in the presence of multiple Y chromosomes may be harmful (Jiang et al., 2010; Lemos et al., 2010; Raznahan et al., 2017).

Polysomic X SCA karyotypes have not been systematically and quantitatively studied by imaging and single cell analysis. Using a modified version of HiCTMap that combines RNA FISH and chromosome paint, we demonstrate that the XCI master regulator *Xist* is present on all supernumerary X chromosome(s) in polysomic X karyotypes in the vast majority of cells in the population. The presence of *Xist* is not affected by the number of Y chromosomes or an increase in Y copy number for example in XXYY cells.

Interestingly, we find changes in the size and shape of Xa and Xi in nuclei from different SCA karyotypes, indicative of changes in the overall level of chromatin compaction of the chromosome. We observe no difference in the size or circularity of Xa or Xi chromosomes as a function of Y copy number. In sharp contrast, the addition of X chromosomes lead to a marked decrease of 20-30% in size of the Xa chromosome with increased X copy number. The Xa chromosome is transcriptionally active and is the functional X chromosome in mammalian cells, but as more X chromosomes are introduced, Xa size decreases. This is in line with the observations that expression of X-linked genes has been found to be lower than expected based on the number of X chromosomes present in patients with increased X copy number (Raznahan *et al.*, 2017). The increased X copy number does not solely involve silencing of genes from the additional inactive X chromosome, but their further repression from the single active X-chromosome by unknown mechanisms (Raznahan *et al.*, 2017). Compensating for the increase in X copy number, we find that the Xa chromosome is compacted and becomes similar in size to the Xi. Additionally, although less striking, any supernumerary Xi chromosomes also show a decrease in size with increasing X copy number, and approach a one to one ratio in size at a single cell level and population level (Figure 3, A and B). Also, we find increased chromosome compaction of the Xa and Xi, as measured by chromosome circularity, with increased X copy number. Chromosomes 18, Xi, and Y are smaller in all SCA karyotypes compared to the active X chromosome. This is likely explained by the fact that chromosomes 18, Xi, and Y are largely heterochromatic while the active X is more open and actively transcribed. Interestingly, we find that not only does the circularity of the Xi chromosome(s) increase with increased X copy number, but also that of the Xa chromosome, with an even higher rate than Xi chromosome(s). The structural changes in shape and size of the Xa and Xi point to a mechanism of global compaction by heterochromatinization of the X chromosome possibly as a way to regulate gene expression.

The reason why polysomic X SCA karyotypes are viable is mostly likely due to the inactivation of the supernumerary X chromosomes by XCI (Tsai *et al.*, 2009). In contrast, polysomy of chromosomes of similar size are often not viable probably due to the increased expression of the additional genes from the extra chromosomes (Tsai *et al.*, 2009; Rosenberg and Rosenberg, 2012). In addition, viable trisomies typically involve significantly smaller and gene poor chromosomes such as 18, 21, and Y (Pearson, 2001; Trump, 2010; Rosenberg and Rosenberg, 2012).

XCI is initiated by the spreading of Xist in cis across the future inactive X chromosome and recruits several proteins, such as lamin B receptor (LBR), polycomb repressive complex 1 (PRC1), and histone deacetylases (HDAC3), that initiates the transcriptional silencing and formation of a repressed nuclear compartment (Clemson *et al.*, 1996; Plath *et al.*, 2002; Chaumeil *et al.*, 2006; Zhao *et al.*, 2008; Pinheiro and Heard, 2017). As XCI is established and maintained, the Xi is enriched for repressive chromatin modification including H2AK119Ub by PRC1, H3K27me3 by PRC2, DNA methylation, and incorporation of the histone variant macroH2A (Plath *et al.*, 2003; Silva *et al.*, 2003; Pinheiro and Heard, 2017). Upon full XCI, the Xi is transcriptionally repressed and tethered to the nuclear periphery via LBR and structurally compacted with escaped genes believed to be localized on the outside of the repressed compartment (Avner and Heard, 2001; Plath *et al.*, 2002; Pinheiro and Heard, 2017). We observe a sizeable decrease in chromosome size and an increase in compaction of the Xa with increasing X copy numbers. This effect is also observed in the Xi, but to a smaller extent. These observations suggest significant alterations in chromosome structure to the Xa in polysomic-X karyotypes. It is possible that, given the higher cellular level of *XIST* in cells containing multiple Xis, XIST may sporadically associate with the Xa and inducing chromatin alterations. This seems unlikely since we could not detect *Xist* staining bound to the Xa in polysomic-X karyotypes. It seems more likely that the size reduction of Xa may be due to Xa heterochromatinization that is *Xist* independent. Prior observation of histone H4 acetylation in XXX, XXXXX, XXXXY, and XXXY karyotypes of the X chromosome showed alterations in deacetylation of histone H4 once the inactive state was established in polyploid-X karyotypes (Leal *et al.*, 1998). In addition, demethylation of histone H4 lysine 20 to monomethyl by Dpy-21 is enhanced on the Xi chromosome in *C. elegans* (Brejc *et al.*, 2017). Furthermore, many of the genes that escape inactivation in SCA karyotypes are upregulated in polyploid X karyotypes are chromatin remodelers (Raznahan *et al.*, 2017), allowing for the possibility that as X copy number increases, repressive chromatin remodelers are deposited on the Xa chromosome, leading to the overall higher order structural changes we observe in the Xa and Xi chromosomes in SCA karyotypes.

Chromosomes are non-randomly arranged in the mammalian cell nucleus (Dundr and Misteli, 2001; Parada and Misteli, 2002; Parada *et al.*, 2004a; Bickmore, 2013). We find no change in position of autosome 18 and sex chromosomes X and Y with increasing X or Y copy number. As previously demonstrated (Belmont *et al.*, 1986; Nagele *et al.*, 1999; Jowhar *et al.*, 2018b) and in these experiments, the Y chromosome has a central position whereas chromosome 18 is equally distributed across the nucleus in all SCA karyotypes. We observe a strong peripheral position of the X chromosome, which becomes stronger with increasing X copy number. However, when we distinguish the X chromosome by XCI status, we observe Xa is more internally positioned compared to Xi and does not show variability between karyotypes. Despite these differences, we observe that the Xa chromosomes of SCA karyotypes are similarly positioned, and the position of Xi chromosomes of SCA karyotypes, despite increasing Xi chromosomes, are unchanged. This suggests that regardless of X copy number, the previous observation of *Xist* mediated Xi tethering to the nuclear lamina is maintained (Chen *et al.*, 2016). Therefore, the spatial positioning of chromosomes remains consistent despite increase in the number of sex chromosomes X or Y.

Taken together, while previous studies have focused on clinical and neurological implications of SCA, we have here probed the effect of supernumeracy of sex chromosomes on genome organization. Our findings document specific nuclear changes in chromosome size and structure, particularly of the X chromosome. These alterations may reflect changes in the global activity of the affected chromosomes and it will be interesting to explore their relationship to gene dosage, most importantly of X-linked genes which may escape XCl, and in this way, contribute to the pathology of SCA diseases.

## Materials and Methods

### Cell Culture

Patient derived primary skin fibroblasts (3 of each karyotype: XO, XX, XXX, XXXX, XXXXY, XXY, XXYY, XY, XYY) were grown in DMEM media with 20% FBS, 2 mM glutamax, and penicillin/streptomycin at 37 °C and 5% CO_2_. Cells were split 1:3 every 3-4 days and kept at a low passage (P3-9). Isolation and expansion of patient derived primary skin fibroblasts was previously described (Jowhar et al. 2018). Normal karyotype was verified by SKY. Patients were enrolled through a phenotypic characterization study of SCA at the National Institute of Mental Health (NIMH) Intramural Research Program. Informed consent and assent was attained from all participants and their parents: all study procedures were approved by an NIH Institutional Review Board.

### High-Throughput Chromosome Paint FISH in 384-Well Plates

High-throughput chromosome paint fluorescence in-situ hybridization (FISH) was performed in triplicate wells as described (Jowhar et al., 2018). Briefly, cells were plated in 384-well CellCarrier Ultra plates at a concentration of 120 cells/ μL (~3,000 cells/well) and grown for 24 hrs. Cells were then fixed in 4% PFA in PBS for 10 min, permeabilized in 0.5% Triton X-100/PBS for 20 min at RT and incubated in 0.1 N HCl for 10 min at RT. Cells were kept in 50% formamide/2X SSC for at least 30 min at RT.

Whole chromosome paint probes of human chromosomes 18, X, and Y were generated in house as previously described (Bischoff *et al.*, 1993; Macville *et al.*, 1997), and labeled using Spectrum Orange (Abbott Molecular), Dy505 (Dyomics), and Dy651 (Dyomics).

For hybridization, a probe mix solution containing 300 ng of each fluorescently labeled chromosome paint probes, 1 μg human COT1 DNA (Sigma-Aldrich Roche), and 30 μg salmon sperm DNA (Ambion) was ethanol precipitated and re-suspended in 12 μL of hybridization buffer (20% dextran sulfate, 50% deionized formamide, 2X SSC, pH 7). The probe mix was then manually added to each well, denatured together with cells at 85°C for 7 min and left to hybridize at 37°C overnight. Excess probe was washed three times with each: 2X SSC at 42°C, 1X SSC and 0.1X SSC at 60°C for 5 min. Cells were finally stained with DAPI in PBS (5 ng/μl) before imaging.

### High-Throughput Chromosome Paint and RNA FISH in 384-Well Plates

A DesignReady *Xist* RNA probe designed by Stellaris (Stellaris, 25µM concentration) was diluted 1:10 in TE buffer, pH 8 and diluted again 1:17 in RNA hybridization buffer (10% formamide, 10% dextran sulfate in 2XSSC) to a final concentration of 150 nM. 12 uL of RNA in hybridization buffer was added to each well and plates were hybridized at 37°C in a humidified chamber for 4 hours. Plates were then washed 3 times for 5 min with 2X SSC at 37°C. Cells were finally stained with DAPI in PBS (5 ng/μl) before imaging. After RNA FISH imaging, DNA FISH and imaging were completed sequentially as described above.

A sub-pixel, intensity based image registration algorithm (Thevenaz *et al.*, 1998) was used to align the maximum intensity projections of the RNA FISH and DNA FISH images. The DAPI channel from the two sequential acquisition sessions was used to estimate rigid body (translation and rotation) transformation which was subsequently applied to the Xist RNA FISH image. The registration parameters were estimated for each field within a well using the ImageJ’s MultiStackRegistration plugin (version 1.46.2) embedded into KNIME image analysis workflow via the ImageJ Macro node.

### High-Throughput Immunofluorescence in 384-Well Plates

Patient derived skin fibroblasts were seeded in 384-well plates 24 hours before fixation. Cells were fixed by the addition of paraformaldehyde to a final concentration of 4% for 10 min, washed once with PBS, permeabilized with PBS/0.5% Triton-X 100 for 10 min, then washed three times with PBS. Immunofluorescence staining was done by 1 hour incubation with primary antibody diluted in block buffer (5% BSA/PBS), wash 3X with PBS, 1 hour incubation with fluorescently labeled secondary antibody and DAPI (2.5 μg/ml) diluted in block buffer (5% BSA/PBS), washed 3X with PBS. The following primary and secondary antibodies were used: mouse-anti-human Ki-67 (BD Transduction Laboratories, 1:500) and Alexa Fluor rabbit-anti-mouse 488 (Invitrogen, 1:1000).

### High-Throughput Cell Cycle Imaging

Cell cycle distribution was measured by high-throughput imaging as previously described (Roukos *et al.*, 2015; Zane *et al.*, 2017). Patient derived skin fibroblasts were seeded in 384-well plates 24 hours before fixation. Cells were fixed by the addition of paraformaldehyde to a final concentration of 4% for 10 min, washed once with PBS, permeabilized with PBS/0.5% Triton-X 100 for 20 min, then washed three times with PBS. DNA was counterstained with 10µg/mL DAPI for 10 min at RT and then washed 3X with PBS. Cells were imaged using the 40X objective in epifluorescence using the Yokogawa CV7000 system. The analysis was conducted as described (Zane et al. 2017).

### Image Acquisition

Cells were imaged in four channels (405, 488, 561, 640 nm excitation lasers) on the Yokogawa CV7000 confocal high-throughput imaging system using a 40X air objective lens (NA 0.95) and 16-bit sCMOS cameras and with no pixel binning. Image Z-stacks of 9-12 images at steps of 0.5 μm were acquired. Under these imaging conditions, the x-y pixel size was 162.5 nm and the field of view size was 416 x 351 µm (2560 x 2160 pixels). At least 12-16 randomly sampled fields were imaged per well.

#### U-Net for Detecting Nuclei and Chromosome Territories

The nucleus segmentation module from our previous work [Jowhar et. al. 2018] was replaced with a fully convolutional neural network, FCN, (Long *et al.*, 2015). The FCN is based on U-Net architecture, a robust semantic segmentation model of detecting objects from biomedical images (Ronneberger *et al.*, 2015). Briefly, the U-Net architecture is composed of an encoder and a decoder component with skip connections from encoder to decoder. We have previously used this architecture successfully for detecting spot-like structures (DNA FISH) in fluorescent images, SpotLearn, (Gudla *et al.*, 2017). The U-Net model for nucleus segmentation is identical to SpotLearn, except for the four spatial down-sampling layers in the encoder section of the network, as well as the four complimentary spatial up-sampling (atrous deconvolution) layers in the decoder section. This modification results in a deeper model and more trainable parameters (7,759,521) than SpotLearn. The second column in the Supplementary Table ST1 describes relevant details about the U-Net model for nucleus segmentation. Since U-Net is a supervised, semantic segmentation model, it requires an input greyscale image and a corresponding binary mask of the objects in the greyscale image. So, to generate the training data for nucleus segmentation, we first selected the maximum intensity projections of the DAPI channel from 12 randomly selected fields of view (FOV, 2560 × 2160 pixels) from an optimization plate. Next, the nuclei in these images were segmented using the seeded watershed and ultrametric contour map as described in our previous work (Jowhar *et al.*, 2018b). Then, we interactively merged all instances of over-segmented nuclei in the FOV into one object. The objects were merged using the “Interactive Label Editor” node in KNIME. Note, we did not correct for under-segmentation (i.e., overlapping or touching) of nuclei. The trained Random Forest classifier in our KNIME analysis workflow will filter these objects, based on nucleus morphology features. This strategy of using existing segmentation method(s) allowed us to rapidly generate binary masks for all DAPI stained nuclei in FOVs.

Prior to training the U-Net model for nucleus detection, we normalized the grey-scale intensities of the DAPI channel to bit-depth of the sCOMS camera(s) on the high throughput microscope (16-bit, 65536 intensity units). After normalization, we extracted 960 patches of 256×256 pixels from the 12 FOVs of the DAPI channel and the corresponding binary mask. We allowed for overlapping windows during the patch extraction.

The U-Net model for nucleus detection was then trained on 864 (90% of 960) patches of 256×256 pixels, using modified Dice loss as described in the spot detection paper (Gudla *et al.*, 2017), mini-batch ADAM optimizer, batch size of 1, initial learning rate of 1e-5, and maximum of 500 epochs. The model performance was evaluated after each epoch on 96 patches (10% of 960) by using the modified Dice loss between the ground truth and model’s prediction. The converged U-Net model for nucleus segmentation had a training and validation modified Dice loss of 0.99097 and 0.95865, respectively. During the inference stage for nucleus detection, the trained U-Net model was applied to the FOV of the DAPI channel (2560×2160 pixels) after normalizing the pixel values to the camera bit-depth (i.e., dividing each pixel intensity value by 65,536). The output generated by the U-Net model is the same size as the input image, but with each pixel value denoting the probability of belonging to the foreground (nucleus). The probability values in the output range from [0, 1] and we applied a hard-threshold of 0.5 to convert pixels into nuclei-class (pixel value > 0.5) or background-class (pixel value <= 0.5). The foreground objects were then subjected to morphological operation for filling holes in the binary masks.

In this work, we also replaced the previously described methodology for CT segmentation (Jowhar *et al.*, 2018b) – the multiscale wavelet transform and Random Forest classifier responsible for filtering out false positive CT detections – with a separate U-Net model. This U-Net model for CT segmentation is only comprised of three spatial down-sampling blocks in the encoder section of the network as well as three complimentary spatial up-sampling blocks (atrous deconvolution) in the decoder section. The third column in the Supplementary Table ST1 describes these details.

For generating the training data for the U-Net model, we first applied the KNIME workflow for chromosome detection from our previous work on the images from an optimization plate (Jowhar *et al.*, 2018b). Then, we manually selected, at random, a collection of 444 CT images, cropped to their nucleus binary mask, for which the CT segmentation mask(s) were visually accurate. The CT images, cropped to their nucleus binary mask, covered all combinations of chromosome X karyotypes, chromosome 18 or chromosome Y karyotypes that we used in this study.

These 444 grey-scale images of CTs and their ground-truth binary masks were then centered and zero-padded to a uniform size of 128×128 pixels. After zero-padding, the intensity values of each CT image were rescaled to [0, 1] using the min-max normalization technique. We increased the training data to 888 by randomly rotating (90, 180, and 270 degrees) the original set of 444 grey-scale CT images and their corresponding binary masks.

The U-Net model for chromosome detection was then trained on 799 (90% of 888) greyscale CT images of 128×128 pixels using modified Dice loss (Gudla *et al.*, 2017), mini-batch ADAM optimizer, batch size of 2, initial learning rate of 1e-5, and maximum of 3000 epochs with early stopping criteria. The model was evaluated after each epoch on 89 greyscale CT images (10% of 888) by using the modified Dice loss between the ground truth and the model’s prediction. The converged U-Net model (epoch number 358) for detecting chromosome territories segmentation had a training and validation modified Dice loss of 0.96692 and 0.93595, respectively.

For inference on unseen CT images (cropped to the binary mask of the nucleus), we apply the min-max normalization of the intensity values in the CT grey-scale image. Then, we center and zero-pad the normalized CT grey-scale image. The centering and zero-padding to a uniform size allows to use the U-Net model to predict on large batches of nuclei (e.g., 256 nuclei). The exact number of nuclei in a batch that can be processed depends on the GPU memory and the complexity of the U-Net model.

The output generated by the U-Net model has the same spatial dimensions as the input, but with pixel values denoting the probability of belonging to the foreground (chromosome territories). The probability values range from [0, 1], and we applied a hard-threshold of 0.99 to convert pixels into CT-class (pixel value >= 0.99) or background-class (pixel value < 0.99). The binary masks of the foreground objects (CT-class) were then labeled into individual objects using the connected-component labeling technique. Lastly, we discarded any CT object of less than 50 pixels for chromosomes X, Y, and 18, and of less than 5 pixels for RNA Xist.

Both the U-Net models were implemented in Python (version 2.7.12) using Keras frontend (version 2.0.5), Tensorflow backend (GPU version, 1.1.0), CUDA library (version 8.0) and cuDNN library (version 5.0). The bespoke python scripts for training and inference were embedded into KNIME (64-bit, Linux, Version 3.2.1) workflow(s) using the “Python Scripting” node. The training and inference of the U-Net model(s) were performed on Biowulf compute node(s) (High Performance Computing Cluster, National Institutes of Health) with Ubuntu 16.04 LTS (64-bit) and equipped with four Kepler 80 NVIDIA GPUs, each with 12 GB graphics memory.

### Sample Numbers and Statistical Analysis

Statistical analyses and graphics were generated using R version 3.3.2., 64-bit with the ggplot2 graphics package (version 2.2.1.) and GraphPad Prism 7 version 7.01. P-values were calculated using one-way ANOVA as indicated in figure legends or two-way ANOVA, and p-values of less than 0.05 were considered statistically significant. All comparisons of karyotypes were made to XX or XY, i.e. XX vs XO, XXX, XXXX, XXXXY, XXY, XXYY, XY, XYY and XY vs XO, XXXXY, XXY, XXYY, XYY. Population values were calculated on a per karyotype and chromosome basis. Replicates were based on three patients, with each experiment containing three technical replicate wells performed in duplicate (N = 6). For each graph, a total of eighteen wells were analyzed per karyotype. Approximately 1,000 nuclei were analyzed per karyotype per experiment.

### Data and Software Availability

KNIME workflows for image analysis, deep learning model definitions along with training weights, and R scripts are available on Github at https://github.com/CBIIT/Misteli-Lab-CCR-NCI/tree/master/Jowhar_SCA_2018

## Acknowledgements

The authors thank the CBIIT Server Team, NCI, National Institutes of Health (NIH) and the Biowulf Cluster, High-Performance Computing Group, CIT, NIH for computational support. This research was supported by funding from the Intramural Research Program of the National Institutes of Health (NIH), National Cancer Institute, and Center for Cancer Research.

## Abbreviations

SCAs: Sex chromosome aneuploidies
DAPI: 4′,6-diamidino-2-phenylindole
HiCTMap: High-throughput chromosome territory mapping
CTs: Chromosome territories
XCI: X chromosome inactivation
Xa: Active X chromosome
Xi: Inactive X chromosome

